# RNA-sequencing reveals strong predominance of *THRA* splicing isoform 2 in the developing and adult human brain

**DOI:** 10.1101/2023.12.22.573013

**Authors:** Eugenio Graceffo, Robert Opitz, Matthias Megges, Heiko Krude, Markus Schuelke

## Abstract

Thyroid hormone receptor alpha (THR*α*) is a nuclear hormone receptor that binds triiodothyronine (T3) and acts as an important transcription factor in development, metabolism and reproduction. THR*α* has in mammals two major splicing isoforms, THR*α*1 and THR*α*2. The better characterized isoform, THR*α*1, is a transcriptional stimulator of genes involved in cell metabolism and growth. The less well characterized isoform, THR*α*2, lacks the Ligand Binding Domain (LBD) and is thought to act as an inhibitor of THRα1 action. The ratio of THR*α*1 to THR*α*2 splicing isoforms is therefore critical for transcriptional regulation in different tissues and during development. However, the expression patterns of both isoforms have not been studied in healthy human tissues or in the developing brain. Given the lack of commercially available isoform-specific antibodies, we addressed this question by analyzing four bulk RNA-sequencing datasets and two scRNA-sequencing datasets to determine the RNA expression levels of human *THRA1* and *THRA2* transcripts in healthy adult tissues and in the developing brain. We demonstrate how 10X Chromium scRNA-seq datasets can be used to perform splicing-sensitive analyses of isoforms that differ at the 3’-end. In all datasets, we discovered a strong predominance of *THRA2* transcripts at all investigated stages of human brain development and in the central nervous system from healthy human adults.

## Introduction

Thyroid hormone receptor alpha (THRα) is a nuclear hormone receptor for triiodothyronine (T3) and is a key factor in many physiological functions throughout the animal kingdom. It acts as a transcription factor for a variety of target genes that are involved in development, metabolism, and growth.^1^ In mammals, two major isoforms are produced by alternative splicing: THRα1 (encoded by *THRA1*), which contains a T3-sensitive ligand-binding domain (LBD), and THRα2 (encoded by *THRA2*), which lacks this T3-binding site and whose activity is thus independent of T3 (**Fig. 1a**).^2^ In the absence of T3, THRα1 binds to the THR-responsive elements (TREs) located in the promoter regions of its target genes and represses gene expression.^3^ Upon T3 binding to THRα1, conformational changes in the receptor promote coactivator binding and ultimate target gene transcription. In contrast, THRα2 is unable to bind T3 and has been shown to act as a weak dominant-negative inhibitor of THRα1 action, possibly *via* protein-protein interaction.^4^ Because of these opposing effects, the molar THRα1 to THRα2 ratio determines the response of the local target cell to T3; the more THRα1 is present, the more T3 response and target gene activation can be expected and conversely, the more THRα2 present, the less target gene expression can be expected (**Fig. 1b**).

**Figure 1:**
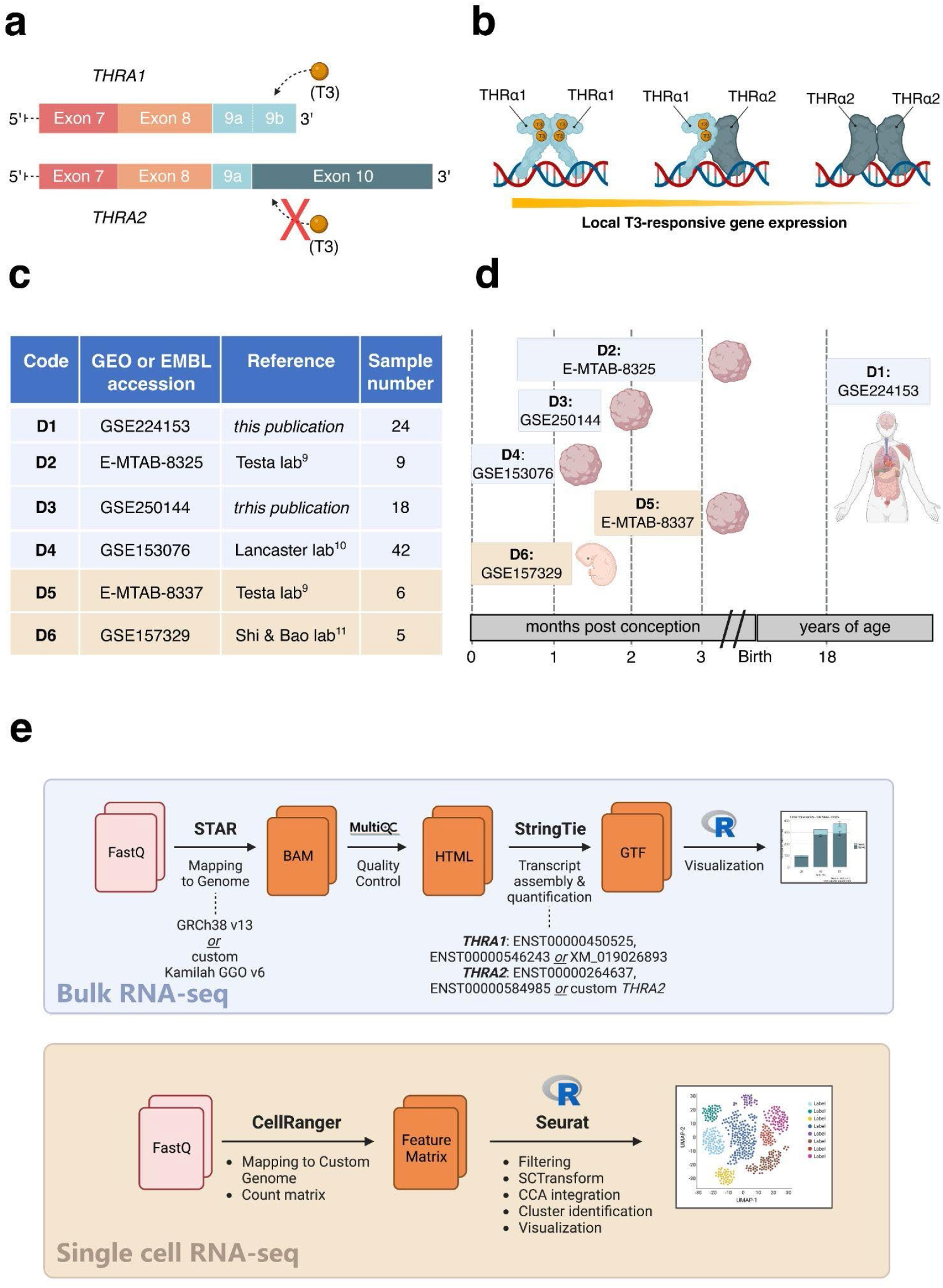
Study overview. **(a)** Schematic representation of the 3’-ends of the *THRA* isoform 1 and 2 mRNA that encode THR*α*1 and THR*α*2, respectively. The orange spheres represent the T3-ligand, and the solid rectangles represent the exons. The schematic highlights that T3 can bind to THR*α*1 but not to THR*α*2. **(b)** Schematic representation of local T3-responsive gene expression based on the abundance of THR*α*1 and THR*α*2. In the presence of the same amount of local T3, cell types that synthesize more THR*α*1 will have more T3-responsive gene expression compared to cell types that synthesize more THR*α*2. **(c)** Overview of the datasets used in this study. Datasets with **blue** background are bulk RNA-seq datasets, while datasets with **beige** background are single-cell RNA-seq datasets. **(d)** Schematic representation of the developmental stages covered by the datasets in relation to human development. Datasets with **blue** background are bulk RNA-seq datasets, while datasets with **beige** background are singlecell RNA-seq datasets. **(e)** Bioinformatic pipelines used to analyze the **bulk RNA-seq datasets** and the **single-cell RNA-seq datasets**.

A recent study in mice demonstrated a predominance of the THR*α*1 isoform over the THR*α*2 isoform in cardiac and intestinal tissues and a predominance of the THR*α*2 isoform in neuronal tissues, thus suggesting a tissue specific regulation and a physiological role for the THR*α*1 to THR*α*2 ratio.^8^

The functional relevance of these differences in T3-response is highlighted by mutations in the coding sequences of the human *THRA* gene.^5,6^ Most of these mutations abolish T3-binding to THR*α*1, thereby increasing the T3-unresponsive receptor pool of the target cell. This results in T3-resistance with a severe phenotype of congenital non-goitrous hypothyroidism (OMIM #614450) with developmental delay, intellectual disability, growth deficit, bradycardia, delayed ossification, and a generally reduced metabolic rate.^5–7^ Despite this proposed physiological relevance, to date, the cellular *THRA1* to *THRA2* ratio has not been systematically investigated during human development or in healthy adults.

To address this knowledge gap, we collected and independently re-analyzed 4 (n=2 publicly available and n=2 in-house generated) human bulk RNA-seq datasets and n=2 publicly available human single-cell RNA-seq datasets generated from healthy adult tissues, brain cortical organoids, and human embryos during early development (**Fig. 1c**). A schematic representation of the developmental stages covered by these datasets in relation to human development is shown in **Fig. 1d**. We found widely varying absolute and relative abundances of the two splicing isoforms in different tissues and, similar to previous studies in mice, a large overrepresentation of the *THRA2* splicing isoform in all neuronal tissues analyzed.

## Results

### *THRA* isoform detection in bulk RNA-seq datasets

We analyzed the expression patterns of human *THRA* splicing isoforms in four different shortread bulk RNA-seq datasets: **D1**, an in-house generated dataset of 24 healthy pooled adult human tissues (GEO database: GSE224153); **D2**, a publicly available dataset of human cortical organoids from the laboratory of Giuseppe Testa (EMBL-EBI database: E-MTAB-8325);^9^ **D3**, an in-house generated dataset of T3-treated (GEO database: GSE250143) and untreated (GEO database: GSE250142) human cortical organoids (GEO database unifying code: GSE250144); **D4**, a publicly available dataset of human and gorilla cortical organoids from the laboratory of Madeline A. Lancaster (GEO database: GSE153076)^10^ (**Fig. 1c** in light blue). We used StringTie to extract the transcripts per million (TPM) values of the two different splicing isoforms (**Fig. 1e**, light blue). The TPM values of *THRA1* and *THRA2* in all bulk RNA-seq datasets are shown in **Figs. 2-5**. In adult tissues, all the nervous system samples expressed high levels of total *THRA* (**Fig. 2**; dataset D1, >100 TPM, full length of the bar). In addition, all nervous system tissues expressed higher levels of *THRA2* compared to *THRA1* (>70% of total *THRA*, dark blue in **Fig. 2**). In the other non-neuronal tissues, with the exception of smooth muscle, *THRA* expression was lower than in neuronal tissues with clear a *THRA2* predominance in thyroid, kidney adrenal, heart, and pancreas while in adipose tissue, skeletal muscle, lung, and colon the *THRA1* isoform (samples below the red line, >50% of total *THRA*, light blue in **Fig. 2**) was more prevalent.

**Figure 2:**
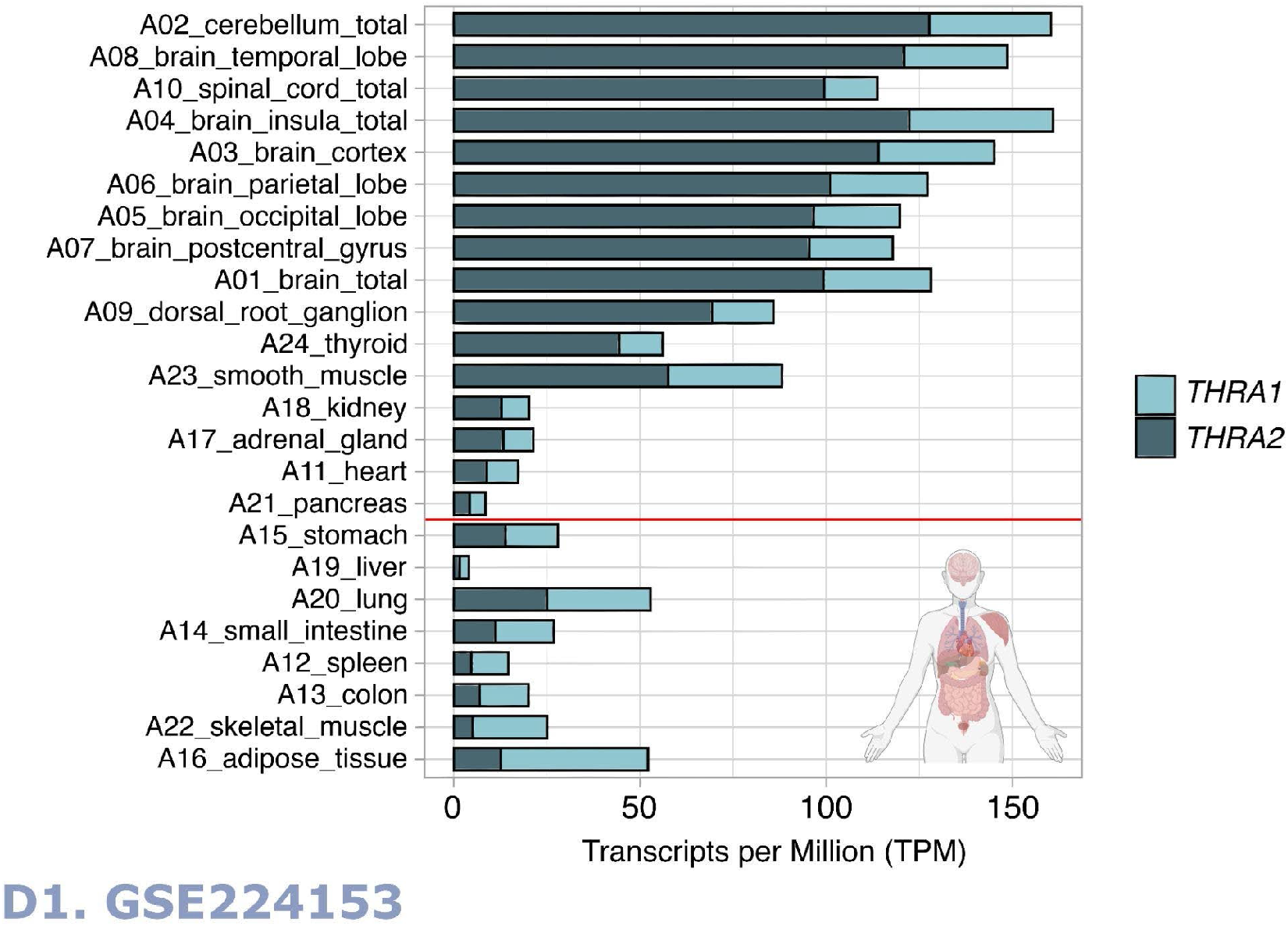
*THRA* isoform expression pattern of dataset D1 GSE224153 in transcripts per million (TPM). The graph shows that all nervous system samples expressed high levels of total *THRA* (full bar length) with a predominance of the *THRA2* isoform (dark blue). Samples are sorted in decreasing order based on the difference between *THRA2* and *THRA1* (light blue). Samples above the red line express more *THRA2* compared to *THRA1*.

In all further analysis we focused on neurons with cortical organoid datasets that allowed us to study developmental changes in *THRA* isoform expression. In all human organoid datasets, we detected increases in total *THRA* mRNA expression over time as well as a strong predominance of *THRA2* over *THRA1* at all time points analyzed (>80% of total *THRA*, dark blue in **Figs. 3a** and **4a**). Notably, this pattern was fully conserved in gorilla cerebral organoids (dataset **D4, Fig. 5**). We also investigated if acute 48 hour treatments of human cortical organoids with high T3 concentrations (50 nM) affect *THRA* isoform expression, but did not find any treatment effects on expression levels of total *THRA* mRNA nor on specific splicing isoforms (**Fig. 4b-d**, one-way ANOVA, p>0.05).

**Figure 3:**
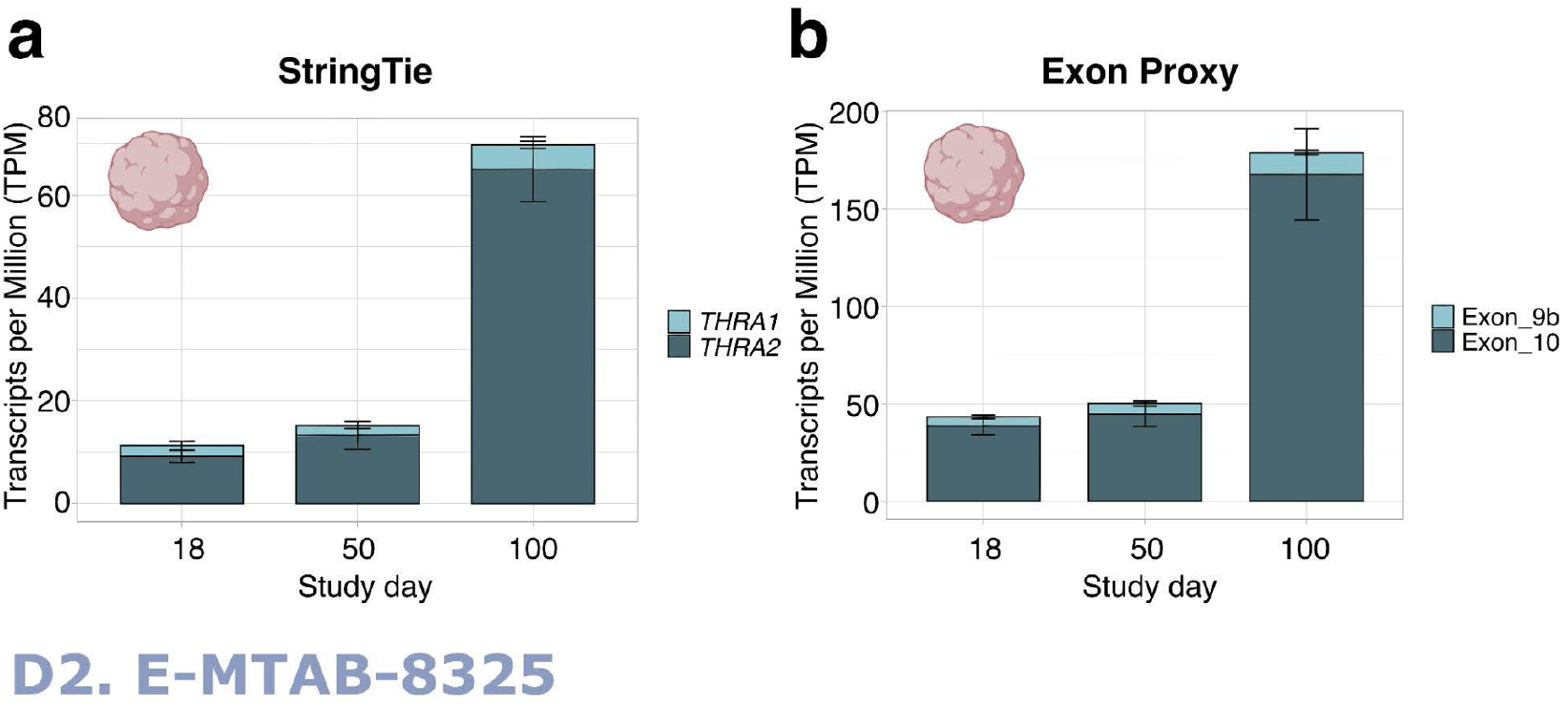
*THRA* isoform expression pattern of dataset D2 E-MTAB-8325 in transcripts per million (TPM). **(a)** TPM calculated with StringTie showing an increase in total *THRA* (full bar length) expression over time and a strong predominance of *THRA2* (dark blue) at all time points. Bar plots show mean ± SEM, n=3 **(b)** TPM calculated using exon 9b as a proxy for *THRA1* (light blue) and exon 10 as a proxy for *THRA2* (dark blue). The plots show a similar expression pattern as in **(a)** indicating that exons 9b and 10 can be used as proxies for isoform expression. Bar plots show mean ± SEM, n=3.

**Figure 4:**
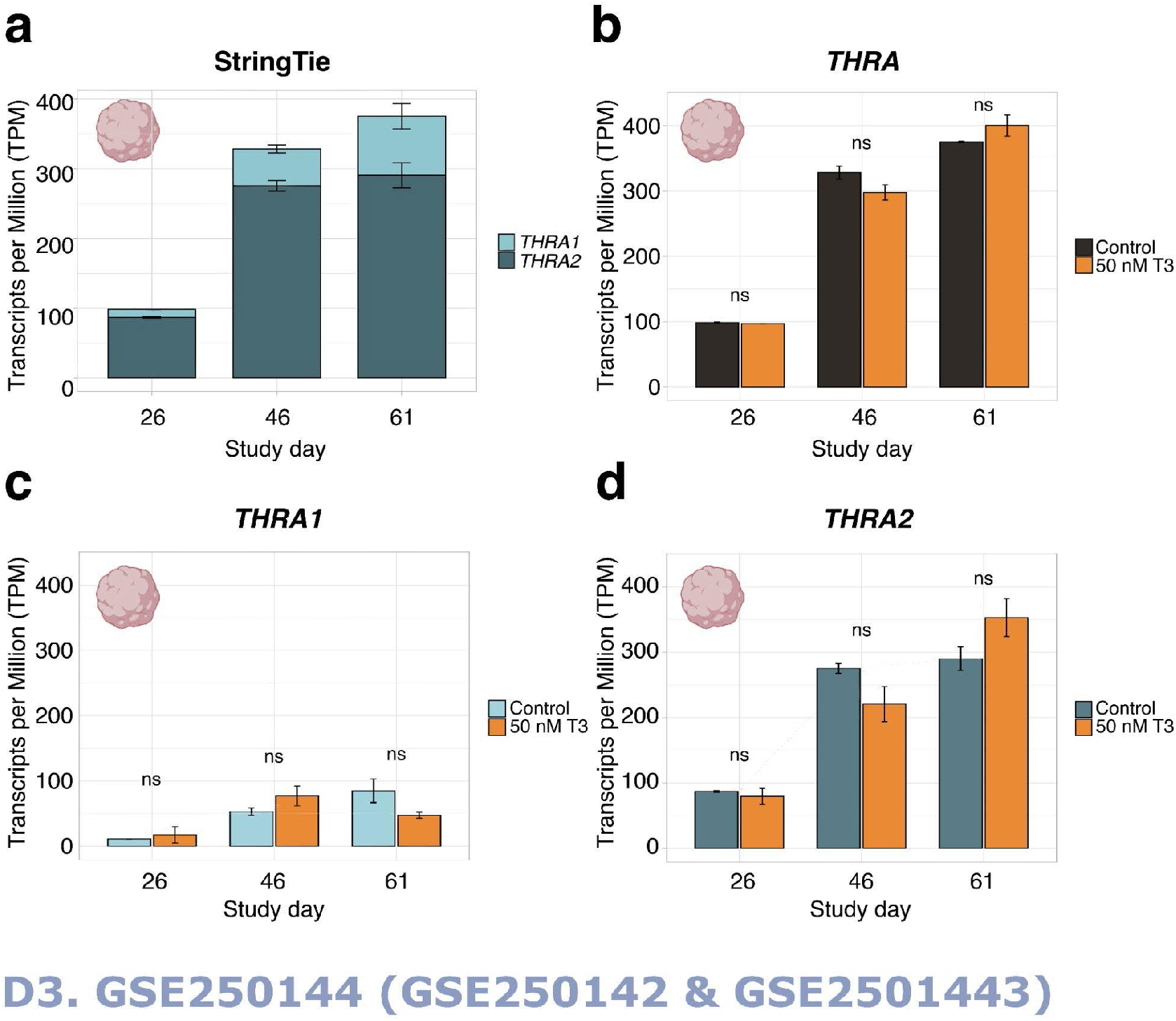
*THRA* isoform expression pattern of dataset D3 GSE250144 in transcripts per million (TPM). **(a)** Graph showing an increase in total *THRA* (full bar length) expression over time and a strong predominance of *THRA2* (dark blue) over *THRA1* (light blue) at all time points. Bar plots show mean ± SEM, n≥2. **(b)** Expression levels of total *THRA* (black) showing no difference between controls and samples treated with 50 nM T3 for 48 h before collection (orange). ns = not significant, mean ± SEM, one-way ANOVA test, n≥2. **(c)** Expression levels of *THRA1* (light blue) showing no difference between controls and samples treated with 50 nM T3 for 48 h before collection (orange). Ns = not significant, error bars depict mean ± SEM, one-way ANOVA test, n≥2. **(d)** Expression levels of *THRA2* (dark blue) showing no difference between controls and samples treated with 50 nM T3 for 48 h before collection (orange). ns = not significant, error bars depict mean ± SEM, one-way ANOVA test, n≥2.

**Figure 5:**
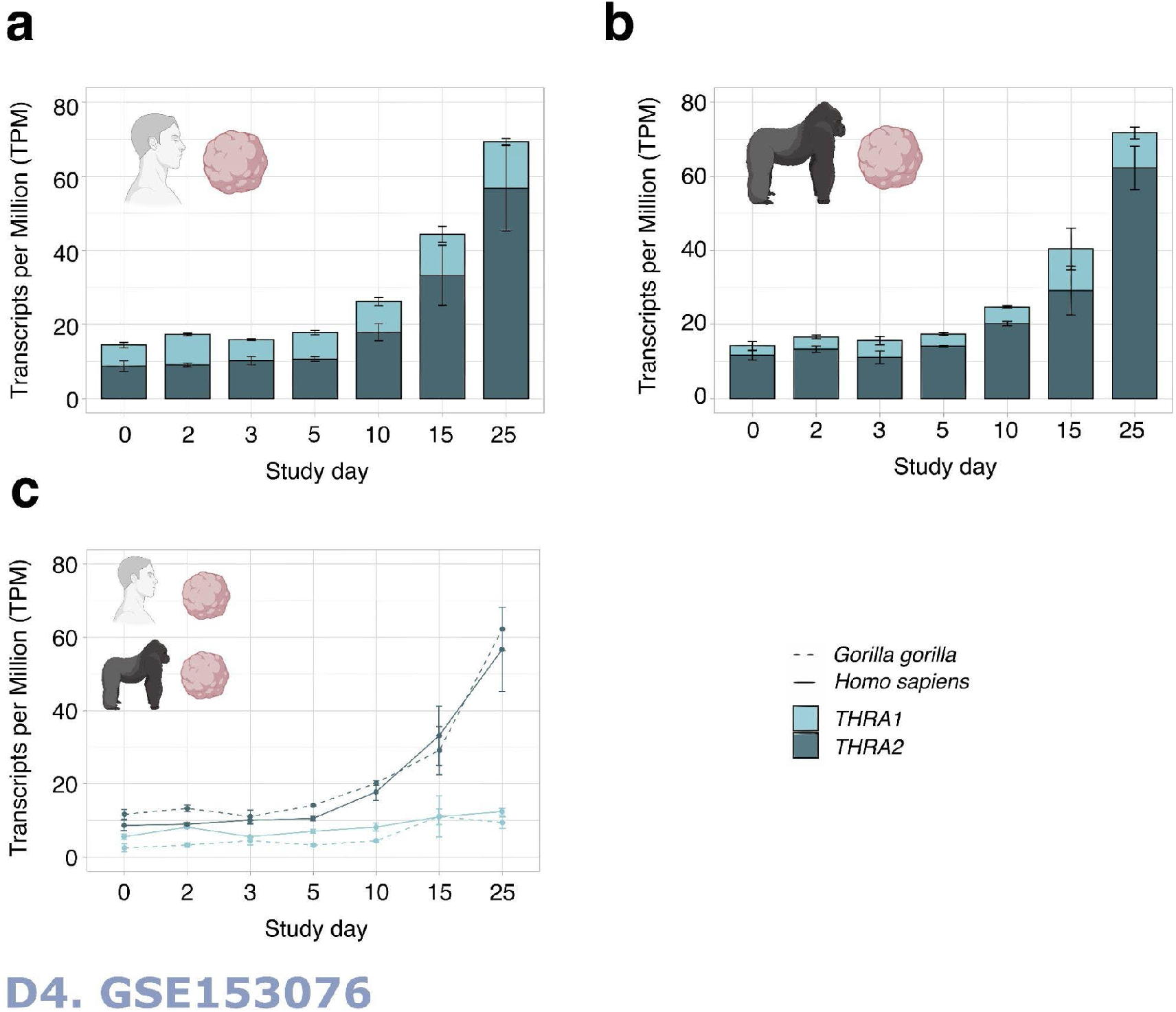
*THRA* isoform expression pattern of dataset D4 GSE153076 in transcripts per million (TPM). **(a)** Graph showing an increase in total *THRA* expression (full bar length) over time and a strong predominance of *THRA2* (dark blue) over *THRA1* (light blue) at all time points in *Homo sapiens* samples. Bar plots show mean ± SEM, n=3. **(b)** Graph showing an increase in total *THRA* expression (total bar length) over time and a strong predominance of *THRA2* (dark blue) over *THRA1* (light blue) at all time points in *Gorilla gorilla* samples. Bar plots show mean ± SEM, n=3. **(c)** Graph comparing the expression levels of *THRA1* (light blue) and *THRA2* (dark blue) in *Homo sapiens* (solid line) *versus Gorilla gorilla* (dashed line), error bars show mean ± SEM, n=3.

### *THRA* splicing isoform detection in single-cell RNA-seq datasets

For the study of *THRA* isoform expression levels in the developing brain at cellular resolution, we mined publicly available single-cell RNA (scRNA)-seq databases. Presently, no scRNA-seq dataset of human cortical organoids or human embryos that had used a splicing isoform-sensitive platform (e.g. Smart-Seq2 and long-read sequencing) is publicly available. In contrast, short-read 10X Chromium datasets are widely available, but tend to have an intrinsic bias towards either the 3’-ends or 5’-ends of their mRNA preparations depending on which specific protocol is used and thus are generally not suitable for isoform-specific analyses. However, the fact that *THRA1* and *THRA2* exactly differ at their 3’-end, provided the justification to re-edit the genome reference to represent *THRA1* exclusively by exon 9b + its 3’UTR and *THRA2* exclusively by exon 10 + its 3’UTR (**Fig. 6a**, see *Methods* section for more details). We validated whether these exons could be used as a proxy to investigate the predominance of either *THRA1* or *THRA2* in bulk RNA-seq dataset **D2**. The relative abundances of *THRA1* and *THRA2* detected with the exon proxy (**Fig. 3b**) are comparable to values detected by StringTie (**Fig. 3a**). Next, we downloaded and bioinformatically re-analyzed two scRNA-seq datasets generated on the 10X Chromium platform: **D5**, a dataset of human cortical organoids from the laboratory of Giuseppe Testa, collected at post neural induction days 50 and 100 (EMBL-EBI database: E-MTAB-8337^9^); and **D6**, a dataset of healthy human embryos from Carnegie stages 12 to 16 (corresponding to post conception days 30-39) from the laboratories of Zhirong Bao and Weiyang Shi (GEO database: GSE157329,^11^ **Fig. 1c** in beige). Cell Ranger v7.1.0 was used to map FastQ files to the custom (GRCh38) human reference genome. We used the Seurat package for data analysis, e.g. to filter low quality cells, normalize the counts, integrate different time points, identify clusters, and visualize results (**Fig. 1e**, beige).

**Figure 6:**
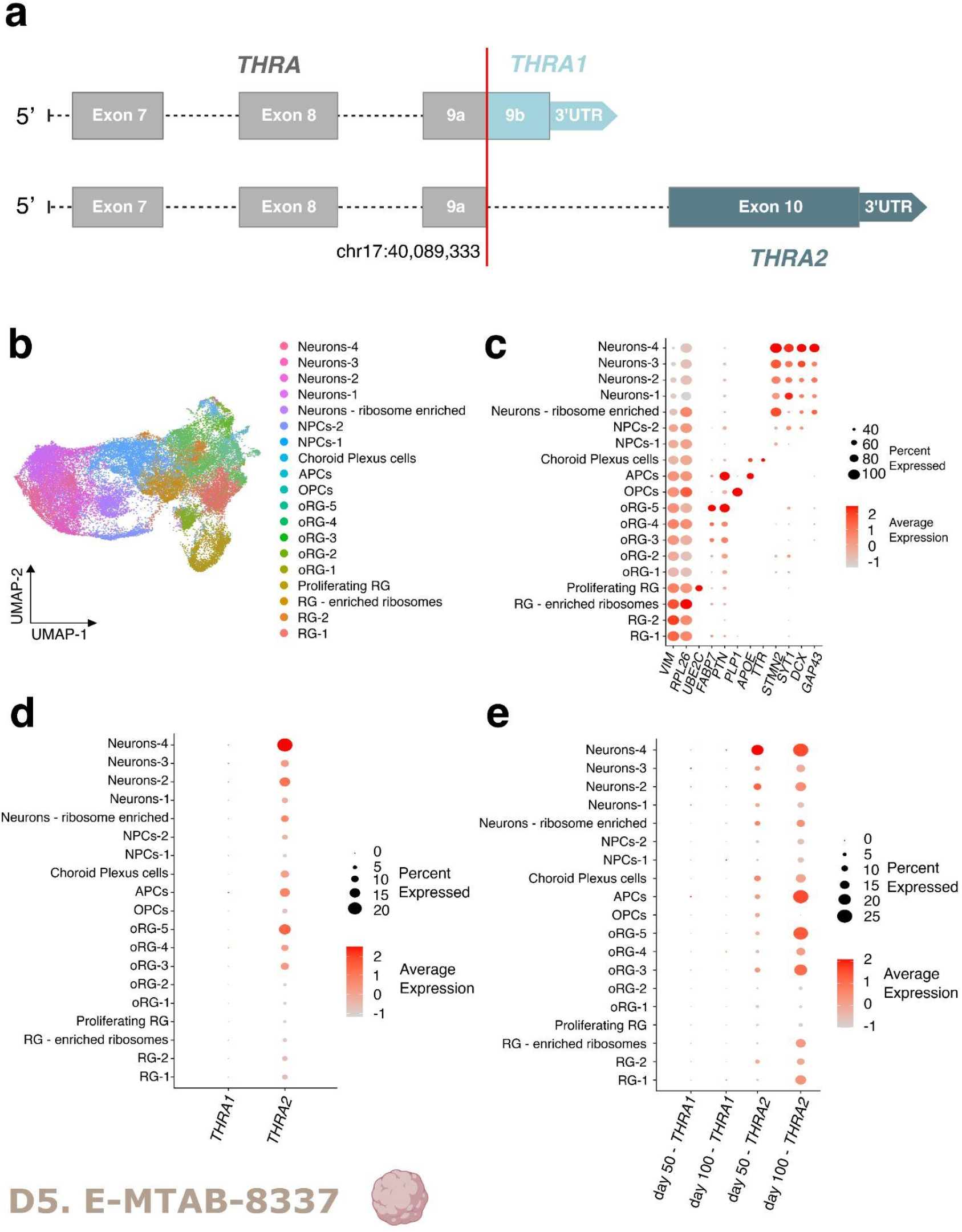
*THRA* isoform expression pattern of human cortical organoids from a single-cell dataset (D5 E-MTAB8337). **(a)** Schematic of edits made to the human reference genome annotation to detect *THRA* isoforms in 10X Chromium scRNA-seq datasets. Exons up to exon 9a (chr17:40,089,33) were considered *THRA* (in gray), exon 9b and its 3’-UTR were mapped to *THRA1* (light blue) specifically and exon 10 and its 3’-UTR were specifically mapped to *THRA2* (dark blue). **(b)** UMAP plot showing the cell types that had been identified by manual curation. **(c)** Dot plot showing the relative expression levels of the gene markers used to identify each population. **Markers:** *IM* for radial glia, *RPL26* for ribosome enrichment, *UBE2C* for proliferating radial glia, *FABP7* and *PTN* for outer radial glia, *PLP1* for the oligodendrocytic lineage, *APOE* for the astrocytic lineage, *TTR* for choroid plexus cells, and *STMN2, SYT1, DCX*, and *GAP43* for neurons **(d)** Dot plot showing the relative expression levels of *THRA1* and *THRA2* isoforms in each cell population (all n=7 samples and time points combined). **(e)** Dot plot showing the relative expression levels of isoforms *THRA1* and *THRA2* in each cell population separated by the time point of analysis. **NPC**, neuronal precursor cell; **APC**, astrocyte precursor cell; **OPC**, oligodendrocyte precursor cell; **oRG**, outer radial glia; **RG**, radial glia; **UMAP**, Uniform Manifold Approximation and Projection.

**Fig. 6b** shows the Uniform Manifold Approximation and Projection (UMAP) representation of dataset **D5**. Identified cell populations recapitulated distinct stages of neuronal differentiation (markers: *GAP43, DCX, SYT1, STMN2*) and of early oligodendrogenesis (marker: *OLP*) and astrocytogenesis (marker: *APOE*) starting from radial glia (marker: *VIM*) and outer radial glia (markers: *FABP7, PTN*). Additionally, we identified *TTR*-positive choroid plexus cells as did the original authors of the dataset. Expression levels of marker genes for each cell cluster are shown in **Fig. 6c**. Expression of *THRA1* and *THRA2* isoforms could be detected across the different cell clusters, while we observed a strong predominance of *THRA2* over *THRA1* in all cell types (**Fig. 6d**). Additionally, when comparing *THRA2* expression levels between cell populations, we detected an increased transcript abundance in more differentiated cell populations compared to immature progenitors. This pattern was confirmed when samples were grouped by the day 50 and day 100 time points (**Fig. 6e**). Interestingly, when comparing expression levels in progenitor populations (i.e. radial glia cells, radial glia cells enriched in ribosomes, and outer radial glia) between day 50 and day 100, we observed higher expression levels of *THRA2* at day 100, indicating that even progenitor cells increase their expression of *THRA2* over time.

**Fig. 7a** shows a UMAP representation of dataset **D6** with cell labels provided by the authors of the dataset. To be able to compare results with dataset D5, we sub-clustered and further analyzed the cell populations originally labeled as “Neuron”, “Neural progenitor”, “Sensory neuron”, and “Schwann cell”. **Fig. 7b** depicts the expression levels of cell markers for each population, namely *DCX, GAP43, STMN2* for neurons, *POU4F1* for sensory neurons, *PLP1, MPZ*, and *SOX10* for Schwann cells, and *PAX6, SOX2*, and *VIM* for neural progenitors. We observed a strong predominance of *THRA2* over *THRA1* in all cell types and a higher *THRA2* expression in differentiated neurons and Schwann cells compared to neural progenitors and sensory neurons (**Fig. 7c**).

**Figure 7:**
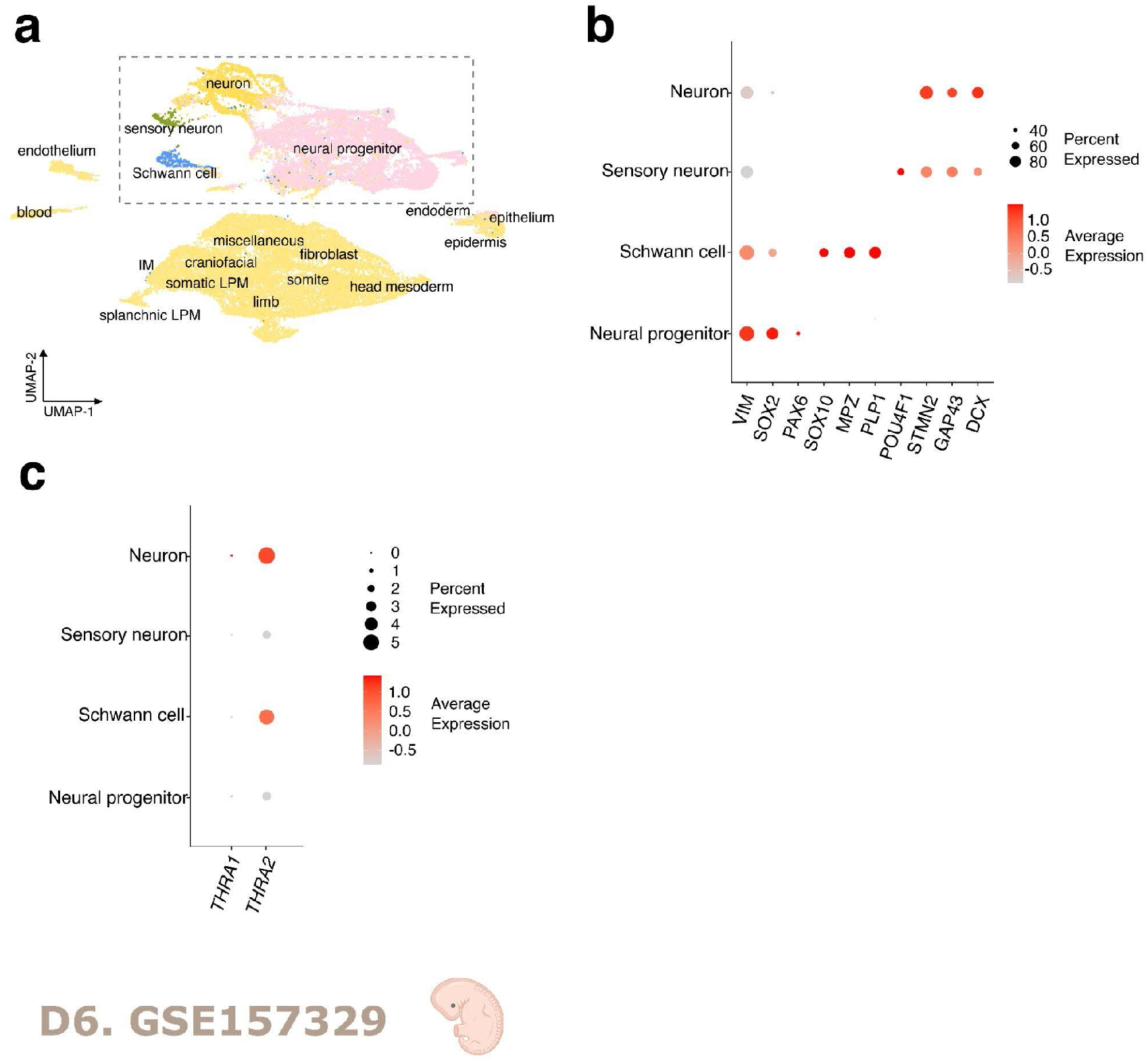
*THRA* isoform expression pattern of human embryos single-cell dataset (D6 GSE157329). **(a)** UMAP plot showing the cell types identified by the authors of the dataset. The dashed rectangle highlights the four cell populations that were sub-clustered and used for further downstream analysis. **(b)** Dot plot showing the relative expression levels of gene markers in the identified cell populations. **Markers:** *VIM, SOX2*, and *PAX6* for neural progenitor cells, *SOX10, MPZ*, and *PLP1* for Schwann cells, *POU4F1* for sensory neurons, and *STMN2, GAP43*, and *DCX* for neurons. **(c)** Dot plot showing the relative expression levels of the *THRA1* and *THRA2* isoforms in each cell population (data derived from all n=6 embryos of Carnegie stages 12-16 combined). **UMAP**, Uniform Manifold Approximation and Projection.

## Discussion

In this study, we independently analyzed four bulk RNA-seq and two single-cell RNA-seq datasets to tease out the expression patterns of the major *THRA* splicing isoforms (*THRA1 versus THRA2*) in healthy adult human tissues and in cortical organoids. We provide an example of how single-cell RNA-seq datasets using the 10X Chromium platform and short-read sequencing, generally thought to be unsuitable for isoform detection, can actually be adapted to distinguish specific isoforms that differ at their 3’-ends by editing the underlying gene annotation file. Based on these datasets, we describe a strong predominance of *THRA2* transcripts at all stages of early human brain development as well as in the central nervous system of healthy adult human tissue. Despite being first described more than 30 years ago, the role of THR*α*2 remains largely unknown.^2,12,13^ Several studies have described its weak dominant-negative effect (DNE) on THR*α*1 mediated gene expression.^14–18^ Authors have suggested that competitive binding of TREs may be the mechanism of action, but more recent findings indicated a lower affinity of THR*α*2 to TREs as compared to THR*α*1.^19^ Therefore, the inhibitory effect of THR*α*2 might be also mediated upstream of receptor binding through its competitive binding of cofactors and the formation of inactive heterodimers.^19^ Thus, THR*α*2 is thought to play a role in modulating the T3-impact on cellular growth and homeostasis. Our study with the demonstration of a strong predominance of THR*α*2 in CNS further extends this idea, and suggests even a protective role of THR*α*2 against thyroid hormone-regulated gene expression at least in the developing brain. This T3-antagonizing role of THR*α*2 might have gained relevance during the evolution of mammals when the fetus falls under the influence of high maternal T3 concentrations.

Whether the detected transcript levels translate into comparable protein levels needs further confirmation. However, given the strong expression of THR*α*2 at the RNA level, one could hypothesize that the *THRA2* transcript may also act as a long non-coding RNA in addition to being a template for protein synthesis. Some examples of such dual-function RNA transcripts have been described in the context of cancer.^20,21^ In addition, considering that exon 10 of *THRA2* is antisense to *NR1D1*, it may down-regulate the expression levels of the resulting protein RevErbɑ which is involved in circadian metabolic control.^22^

In addition, both the expression and the activity of *THRA* isoforms are regulated post-transcriptionally and post-translationally. In particular, dephosphorylation of THR*α*2 has been shown to increase the DNA binding affinity and thus the inhibitory function of THR*α*2^23^ and sumoylation has been shown to affect interaction with other TRs and thus gene expression.^24,25^

Together, our findings with the detected difference in THR*α*1 *versus* THR*α*2 expression should be kept in mind when inferring biological mechanisms from molecular studies that do not distinguish between the two isoforms. Further expression studies should investigate the isoform expression pattern at later stages of fetal brain development and at postnatal life stages.

Finally, open science, data sharing, and reuse have recently advanced scientific progress by promoting collaboration, transparency, and cost-effectiveness. A large number of scRNA-seq datasets generated on the 10X Chromium platform are available for researchers to use. The 10X Chromium platform is known for its high throughput and scalability and is widely used.^26^ However, these datasets are generally not suitable for isoform-sensitive analyses due to the short-read sequencing processing and the 3’-end or 5’-end bias of the library preparation. Given the unavailability of open access single-cell datasets of human cortical organoids generated by isoform-sensitive platforms such as Smart-seq2 and long-read sequencing, we took advantage of the fact that *THRA1* and *THRA2* differ exactly at their 3’-end and edited a reference genome to map *THRA1* and *THRA2* to their specific exons. Despite the relatively shallow sequencing depth of the 10X Chromium platform, both isoforms were detected and the difference in UMI counts between the two isoforms was large enough to confirm the predominance of *THRA2* in all cell populations. This study provides an example of how widely available scRNA-seq datasets generated with the 10X Chromium platform can be used to perform isoform-sensitive analyses in specific scenarios.

## Methods

### Healthy adult human tissues – Dataset D1

Dataset **D1** includes 10 different nervous system tissues and 14 non-neuronal tissues, each sample pooled from an average of six healthy individuals (range 1-21 individuals), male/female, Asian, Caucasian and African American, aged 18-89 years. Causes of death were reported as sudden death or traffic accidents. High-quality total RNA was purchased from Takara (www.takarabio.com). Total RNA was isolated by a guanidinium thiocyanate method.^27^ Total RNA concentration, integrity, and purity were analyzed by capillary electrophoresis (CE) using the Agilent 2100 Bioanalyzer and the Agilent Fragment Analyzer (Agilent Technologies, Santa Clara, CA, USA). A detailed overview of the data set is provided in **Supplementary Table 1**.

### Human cortical organoids – Dataset D3

Cortical organoids were generated as described in a previous publication.^28^ Briefly, cerebral organoids were derived from hiPSC of n=3 healthy donors using the STEMdiff™ Cerebral Organoid Kit (StemCell Technologies, #08570, #08571) according to the manufacturer’s instructions and collected on days 21, 41 and 61. A subset of organoids was treated with 50 nM T3 for 48 hours prior to tissue collection. Samples were snap frozen in liquid nitrogen and stored at -80°C until RNA extraction. RNA was extracted using the NucleoSpin RNA Plus Kit from Macherey and Nagel (#740984) according to the manufacturer’s guidelines. A detailed overview of the data set is provided in **Supplementary Table 2**.

### Bulk RNA sequencing – Datasets D1 and D3

At least 500 ng of each sample from the **D1** and **D3** datasets were sent out for bulk RNAsequencing which was performed at the Beijing Genomics Institute (BGI, Shenzhen/Hong Kong, China, www.genomics.cn) using their DNBseq platform on a strand-specific cDNA library generating 60 million 100 bp paired-end reads. The library preparation included an mRNA enrichment step using oligo(dT)-attached magnetic beads.

### Publicly available datasets – D2, D4, D5, and D6

FASTQ files of datasets **D2, D4, D5**, and **D6** were downloaded from either the Gene Expression Omnibus (GEO) database (**D4** GSE153076, **D6** GSE157329) or the EMBL-EBI database (**D2** E-MTAB-8325, D5: E-MTAB-8337). Specifically, for dataset **D2** we used the paired-end reads from all untreated samples that were used as controls by the original authors (n=3 replicates each from day 18, 50, and 100 post neuronal induction). For dataset **D4** we used all the available samples (n=3 replicates each from days 0, 2,3,5, 10, 15, and 25 post neuronal induction for human cortical organoids and n=3 replicates from days 0, 2, 3, 5, 10, 15, and 25 for gorilla cortical organoids). For dataset **D5**, we downloaded all untreated samples that were used as controls by the authors (n=3 replicates each from days 50 and 100 post neuronal induction). For dataset **D6**, we used all samples containing the head sections of the embryos, namely *embryo1-head_trunk, embryo2-head, Emn*.*04-head, Emb*.*05-head, Emb*.*06-head a*, and *Emb*.*06-head b* (n=72 FASTQ files). Details of all samples included for analysis are summarized in **Supplementary Table 3**.

### Bioinformatic analyses of bulk RNA-seq datasets – D1, D2, D3, D4

Datasets **D1, D2, D3**, and **D4** were analyzed using the same pipeline (**Fig. 1e**, light blue). Sequence quality was assessed using FastQC v0.11.8 (www.bioinformatics.babraham.ac.uk/projects/fastqc) and MultiQC v1.6.^29^ Reads were mapped to the EMBL human genome 38, patch release 13 (**D1, D2, D3**, and **D4** human samples) or to the modified UCSC Kamilah gorilla genome GGO, version 6 (**D4** gorilla samples, see paragraph below for details) using the splice-aware aligner STAR v2.7.10a.^30^ BAM files were sorted and indexed using SAMtools v1.9.^31^ Mapping quality was assessed by calculating `junction_saturation.py`, `inner_distance.py`, `read_distribution.py`, `read_duplication.py` with RSeQC v2.6.4.^32^ StringTie v2.1.7 was used to extract the transcripts per million (TPM) values of all the known *THRA* variants in each tissue.^33^ For *THRA1* expression, we summed up the TPM values of the variants containing exon 9b (human samples: ENST00000450525 and ENST0000054624; gorilla samples: XM_019026893.2). For *THRA2* expression, we summed up the TPM values of the variants containing exon 10 (human samples: ENST00000264637 and ENST00000584985; gorilla samples: custom *THRA2*, see below for details). FeatureCounts v2.0.1 was used to extract read counts for exon 9b (chr17:40,089,334-40,092,627, (+)) and exon 10 (chr17:40,093,020-40,093,867, (+)) for proxy validation.^34^ Data visualization and statistical analysis were performed in R v4.0.0 (https://www.r-project.org/).

### Generation of a custom gorilla genome reference with *THRA2* locus

To date, the mRNA transcript of the gorilla *THRA2* isoform has not been deposited in the NCBI Reference Transcriptome. To localize the genomic coordinates of *THRA* exon 10 in the gorilla genome, we manually BLAST aligned the human *THRA* exon 10 sequence to the gorilla genome (chr5:41,857,943-41,858,305(-)). We then created a custom reference genome by adding an entry for *THRA2* (transcript chr5:41,857,944-41,888,473 (-); exons 1 to 9 same as *THRA1* entry, exon 10 chr5:41,857,944-41,858,305(-); 3’UTR chr5:41,857,457-41,857,944(-); stop codon chr5:41,857,941-41,857,943(-)). The modified annotation of the gorilla reference genome has been uploaded to figshare (https://doi.org/10.6084/m9.figshare.24486601)

### Bioinformatic analyses of single-cell RNA-seq datasets – D5, D6

Datasets **D5** and **D6** were analyzed individually using a similar pipeline (**Fig. 1e**, beige). Sequencing quality was assessed using FastQC v0.11.8 and MultiQC v1.6. Cellranger v7.1.0 was used to map the reads to our custom human reference genome (see paragraph below for more details) and to extract count matrices.^35^ We followed a standard workflow for data preprocessing using the Seurat package v4.3.0^36^ in R v4.0.0.

Quality control preprocessing included filtering out low quality cells with mitochondrial gene count greater than 4-10% and cells with feature counts less than 200 or 1000 and greater than 9,000 or the 90^th^ centile. **Table 1** shows the quality control metrics for each sample (*i*.*e*. number of cells before filtering, median number of genes per cell, % mitochondrial gene threshold, feature count threshold and number of cells after filtering). Data were normalized and scaled (regressing out % mitochondrial genes), and highly variable genes were calculated using the `SCTransform()` function of the Seurat package .

**Table 1:** Filtering thresholds used to analyze each single-cell RNA-seq dataset.

| Dataset Code | Origin | GEO or EMBL accession numbers | Submitted file name | Study day | Number of cells before filtering | Median genes per cell | Threshold % of mitochondrial genes | Threshold number of features | Number of cells after filtering |
| --- | --- | --- | --- | --- | --- | --- | --- | --- | --- |
| D5 | Testa lab | E-MTAB-8337 | 3391B_D50 | 50 | 5,576 | 2,566 | < 5% | >200 & < 9000 | 5,558 |
| D5 | Testa lab | E-MTAB-8337 | kolf2c1day50 | 50 | 11,785 | 2,576 | < 4% | >200 & < 9000 | 11,739 |
| D5 | Testa lab | E-MTAB-8337 | G1_MIF1D50 | 50 | 4,602 | 1,701 | < 6% | >200 & < 9000 | 4,577 |
| D5 | Testa lab | E-MTAB-8337 | S20271_156 | 100 | 2,345 | 2,379 | < 8% | >200 & < 9000 | 2,251 |
| D5 | Testa lab | E-MTAB-8337 | S20269_154 | 100 | 1,389 | 1,853 | < 9% | >200 & < 9000 | 1,356 |
| D5 | Testa lab | E-MTAB-8337 | S20265_136 | 100 | 9,610 | 1,754 | < 10% | >200 & < 9000 | 9,562 |
| D5 | Testa lab | E-MTAB-8337 | S20276_177 | 100 | 3,188 | 1,619 | < 5% | >200 & < 9000 | 3,168 |
| D6 | Shi and Bao lab | GSE157329 | embryo1-head_trunk | CS 12 | 21,570 | 2,879 | < 10% | >1000 & < 90th quantile | 13,462 |
| D6 | Shi and Bao lab | GSE157329 | embryo2-head | CS 13-14 | 13,193 | 1,546 | < 10% | >1000 & < 90th quantile | 4,386 |
| D6 | Shi and Bao lab | GSE157329 | Emb.04-head | CS 13-14 | 18,249 | 2,321 | < 10% | >1000 & < 90th quantile | 8,734 |
| D6 | Shi and Bao lab | GSE157329 | Emb.05-head | CS 15-16 | 17,246 | 2,894 | < 10% | >1000 & < 90th quantile | 12,926 |
| D6 | Shi and Bao lab | GSE157329 | Emb.06-head a | CS 15-16 | 8,109 | 2,382 | < 10% | >1000 & < 90th quantile | 3,406 |
| D6 | Shi and Bao lab | GSE157329 | Emb.06-head b | CS 15-16 | 8,357 | 1,942 | < 10% | >1000 & < 90th quantile | 3,426 |

In both datasets, samples corresponding to different time points and replicates were integrated using the Seurat package and the CCA Pearson residual method. Specifically, we sequentially ran Seurat’s `SelectIntegrationFeatures()`, `PrepSCTIntegration()`, `FindIntegrationAnchors()`, and `IntegrateData()` with 3,000 integration features.

For dataset **D5**, we identified clusters using the `FindClusters()` function (resolution = 0.5) and manually annotated cell types based on the original authors’ annotations and cell type-specific markers found through multiple rounds of sub-clustering. For dataset **D6**, we used the annotations from the original publication and sub-clustered and further analyzed the cell populations originally labeled as “Neuron”, “Neural progenitor”, “Sensory neuron”, and “Schwann cell”.

### Generation of a customized human genome reference for the analysis of the scRNA-seq datasets

We edited the genome reference to map exon 9b and the corresponding UTR, stop codon and CDS exclusively to *THRA1* and exon 10 exclusively to *THRA2* (**Fig. 6a**). To achieve this, entries corresponding to exon 10 (chr17:40,093,020-40,093,613(+), chr17:40,093,137-40,093,867(+) and chr17:40,093,020-40,093,867(+)) were given a new gene_id, transcript_id and gene_name “*THRA2*” to make it count as a separate gene and allow mapping with Cellranger. Entries corresponding to exon 9b were given a new gene_id, transcript_id and gene_name “*THRA1*” and their start position was edited to avoid overlapping with exon 9a mapping to *THRA* (chr17:40,089,334-40,092,627(+) and chr17:40,089,334-40,089,730(+)). Exons 1 to exon 9a have been set to map to *THRA*. The generated custom human reference genome has been uploaded to figshare (https://doi.org/10.6084/m9.figshare.24486535.v2).

## Data Availability

The paired-end FASTQ files of the **D1** dataset have been submitted to the GEne Omnibus (GEO) database under the accession number GSE224153. RNA sample metadata, e.g. Takara lot numbers, and quality statistics are available in the accompanying metadata spreadsheet of the GSE224153 database entry and in **Supplementary Table 1**. The FASTQ files of the **D3** dataset have been submitted to the GEO database under the accession number GSE250142 (native cortical organoids) and GSE250143 (T3-treated cortical organoid) and metadata are available in **Supplementary Table 2** (dataset**s** GSE250142 and GSE250143 are combined into dataset GSE250144). Datasets **D2, D4, D5** and **D6** are publicly available under the accession numbers given above.

## Code Availability

The R codes used to analyze both bulk and single-cell datasets are available at https://github.com/eugeniograceffo/Graceffo_et_al_2023_Nature_Scientific_Data

Mapping of the bulk RNA-seq FASTQ files was done with the following STAR options:

`STAR --runThreadN 60 \`

`--readFilesCommand gunzip -c \`

`--twopassMode Basic \`

`--outFilterIntronMotifs RemoveNoncanonical \`

`--outSAMstrandField intronMotif \`

`--outSAMtype BAM Unsorted \`

`--alignSoftClipAtReferenceEnds No \`

`--outFilterMatchNmin 100 \`

`--outSAMattrIHstart 0 \`

`--alignEndsType EndToEnd`

StringTie was run as follows:

`stringtie -o ${file%.sorted.bam}.stringtie_output.gtf \`

`-e \`

`--rf \`

`-p 60 \`

`-A ${file%.sorted.bam}.gene_abund.tab \`

`-C ${file%.sorted.bam}.cov_refs.gtf`

## Acknowledgements

This research was funded by grants of the Einstein Stiftung Berlin (ECN scholarship to E.G.), of the Deutsche Forschungsgemeinschaft (DFG; German Research Foundation) under Germany’s Excellence Strategy EXC-2049-390688087 to M.S., and by the Transregio Collaborative Research Center “LocoTact” (TRR 296 TP06) to H.K. and M.S.

## Author contributions

E.G. and M.S. conceived the study, R.O. and M.M. generated the cortical organoid samples (dataset **D3**), E.G. performed the RNA extraction experiments, E.G. and M.S. analyzed the bulk RNA-seq data, E.G. analyzed the scRNA-seq data, H.K. and M.S. supervised the research and provided funding. E.G. wrote the first draft of the manuscript. All authors read and edited the manuscript for intellectual content and consented to its publication.

## Competing interests

The authors declare no competing interests.

**Supplementary Table 1:** Metadata of dataset D1 GSE224153.

**Supplementary Table 2:** Metadata of dataset D3 GSE250144 (GSE250142 + GSE250143).

**Supplementary Table 3:** Overview of publicly available datasets used in the analysis.

## Notes

https://doi.org/10.6084/m9.figshare.24486535.v2

https://www.ncbi.nlm.nih.gov/geo/query/acc.cgi?&acc=GSE224153

https://www.ncbi.nlm.nih.gov/geo/query/acc.cgi?acc=GSE250142

https://www.ncbi.nlm.nih.gov/geo/query/acc.cgi?acc=GSE250143

### Competing Interest Statement

The authors have declared no competing interest.

